# Control of gastruloid patterning and morphogenesis by the Erk and Akt signaling pathways

**DOI:** 10.1101/2023.01.27.525895

**Authors:** Evan J. Underhill, Jared E. Toettcher

## Abstract

Fibroblast growth factor (FGF) dependent elongation along an anterior-posterior (A-P) axis is a conserved feature of vertebrate embryogenesis. A-P axis elongation can also be reproduced in 3D cell culture models termed gastruloids, enabling dissection of this process in a controlled, minimal context. Here, we set out to determine how gastruloid posterior elongation depends on the Erk and Akt pathways, canonical downstream effectors of FGF signaling. We find that gastruloids exhibit reproducible posterior-to-anterior gradients in Erk and Akt phosphorylation that are generated independently and correlate with distinct zones of tissue movement, cell proliferation, and expression of cell motility and adhesion regulators. Pharmacological inhibition of FGFR, Erk, or Akt signaling impairs gastruloid elongation, and quantification of signaling gradients reveals how these patterns interact and scale with A-P axis length. Using global inhibitors and activators of each pathway, we find that a gradient of Ras/Erk signaling is required for the establishment of appropriately localized domains of E-cadherin, Snail, and Brachyury expression, whereas perturbing PI3K/Akt signaling alters proliferation but not patterning. Taken together, our data demonstrate that graded PI3K/Akt and Ras/Erk signaling provide spatial information to control proliferation and cell-cell adhesion during gastruloid elongation.

## Introduction

The shaping of a multicellular structure is a physical process that is driven by dynamic fields of cell proliferation, tissue mechanics, and active cellular forces^1^. These dynamic fields are often modulated by gradients in cell signaling, which are either pre-patterned, as in the case of maternally-localized mRNAs in the early *Drosophila* embryo^2,3^, or arise spontaneously, as seen in numerous examples of self-organization throughout development (e.g., digit emergence^4^, hair follicle formation^5^, and lung branching^6^). One particularly well-studied example of morphogenesis is vertebrate anterior-posterior (A-P) axis elongation, during which the posterior-most tailbud domain unidirectionally elongates while maintaining a roughly constant lateral width^7^. Across vertebrate species, this process is thought to be driven by localized cell proliferation in the tail bud as well as a gradient in tissue-scale mechanical properties that effectively fluidizes the posterior domain^8^. Precisely how biochemical signals are integrated to generate these gradients in tissue properties remains unknown.

Embryonic organoids can provide a window into early development in a reduced-complexity context, while also circumventing the experimental challenges of the post-implantation mammalian embryo. The mouse gastruloid, for example, is a model system that can recapitulate many developmental processes, including the formation of the three body axes, Hox gene patterning, tailbud elongation, and even somitogenesis^9–11^. Beginning from a collection of mouse embryonic stem cells (mESCs), the gastruloid spontaneously develops asymmetric gene expression over the course of 4 days, followed by the emergence and rapid outgrowth of a posterior domain resembling the vertebrate tailbud. This axial elongation produces a dramatic morphological change over 24 h, causing gastruloids to transition from an approximately 350 μm diameter spheroid to a 1 mm long, 200 μm wide cylindrical tissue. The quick and reproducible development of gastruloids, coupled with the ease of mESC engineering and perturbation, makes this simplified model system highly amenable to the study of signaling gradients and morphogenesis *in vitro*. Defining the regulatory processes that drive axial elongation in gastruloids could shed light on how similar morphogenetic events are achieved in the embryo; they may also provide a basis for the future engineering of user-defined morphogenesis programs.

Fibroblast growth factor (FGF) signaling plays a central role in vertebrate A-P axis specification and elongation. FGF ligands are produced at the tailbud, establishing a posterior-to-anterior gradient in ligand concentration along the A-P axis^12^. This signal is essential for proper axial development, as mouse embryos with conditionally inactive *Fgf4/8* exhibit a truncated axis and impaired somite formation^13^. Two canonical FGFR effector pathways, the Ras/Erk and PI3K/Akt signaling cascades, also exhibit graded activity along the anteroposterior axis of the late-gastrulation mouse embryo^12,14^. Ras/Erk signaling is a key regulator of somite formation in vertebrate species^15,16^, and has been shown in gastruloids to regulate the speed of the determination front^10,17^ and the maintenance of an axial progenitor population^18^. The role of PI3K/Akt signaling during vertebrate elongation is comparatively understudied: while PI3K/Akt signaling has been implicated in the migration of mesodermal cells during chick and zebrafish gastrulation^19,20^, similar experiments have not been reported in mouse models^21^.

Here, we sought to characterize the role of the Ras/Erk and PI3K/Akt signaling pathways in the growth and morphogenesis of gastruloids derived from mouse embryonic stem cells. We first established that the Erk and Akt pathways form highly reproducible posterior-to-anterior gradients that span ~500 μm, with Erk exhibiting an additional secondary peak of phosphorylation near the gastruloid midpoint. Quantification of morphological and signaling changes in response to pathway inhibitors revealed that both pathways regulate posterior elongation, but with unexpected regulatory complexity: namely, the primary input to Akt signaling is FGFR-independent, and the secondary Erk peak scales proportionally with overall gastruloid length. Finally, we find that local Erk and Akt activity regulate axial elongation *via* distinct downstream cellular processes: both Erk and Akt regulate total cell number and overall gastruloid size, whereas the Erk gradient is also required for the proper expression and spatial patterning of genes including T/Brachyury, E-cadherin, and Snail. Together, our data indicate that independent spatial patterns of Erk and Akt activity each play distinct but essential functional roles in gastruloid axial elongation.

## Results

### Elongating mouse gastruloids possess posterior-to-anterior patterns of Erk and Akt activity

We first set out to quantify the spatial patterns of Erk and Akt activity during gastruloid elongation. We generated gastruloids using a Bra^GFP^ mouse embryonic stem cell line^22^ in which GFP expression is driven from one of the endogenous T/Brachyury (Bra) loci. Bra marks mesodermal cells in the posterior domain, providing a useful fiduciary pattern against which to compare the spatial distributions of additional signaling pathways and target genes. We grew gastruloids using a standard protocol after sorting exactly 200 mESCs into each well of a round-bottomed plate to reduce well-to-well variability (**Figure 1A**; **Methods**). This protocol resulted in aggregates with a major axis of approximately 1 mm at the end of Day 5 (120 h); a small number (fewer than 5%) of the aggregates failed to elongate or formed multiple posterior domains. At the end of Day 5 we fixed and stained gastruloids for either doubly phosphorylated Erk (ppErk) or phosphorylated Akt (pAkt), marking the active forms of both kinases. We then imaged the spatial distribution of the phosphorylated kinases, Bra^GFP^, and cell nuclei using DAPI. These experiments revealed that both ppErk and pAkt formed strong posterior-to-anterior signaling gradients, consistent with observations of differential activity from these pathways in the mouse embryo^12,14^ (**Figure 1B**). We observed substantial cell-to-cell variability in ppErk staining at the posterior pole, with comparatively low heterogeneity in the pAkt pattern (**Figure 1B**; **Figure S1A**).

**Figure 1:**
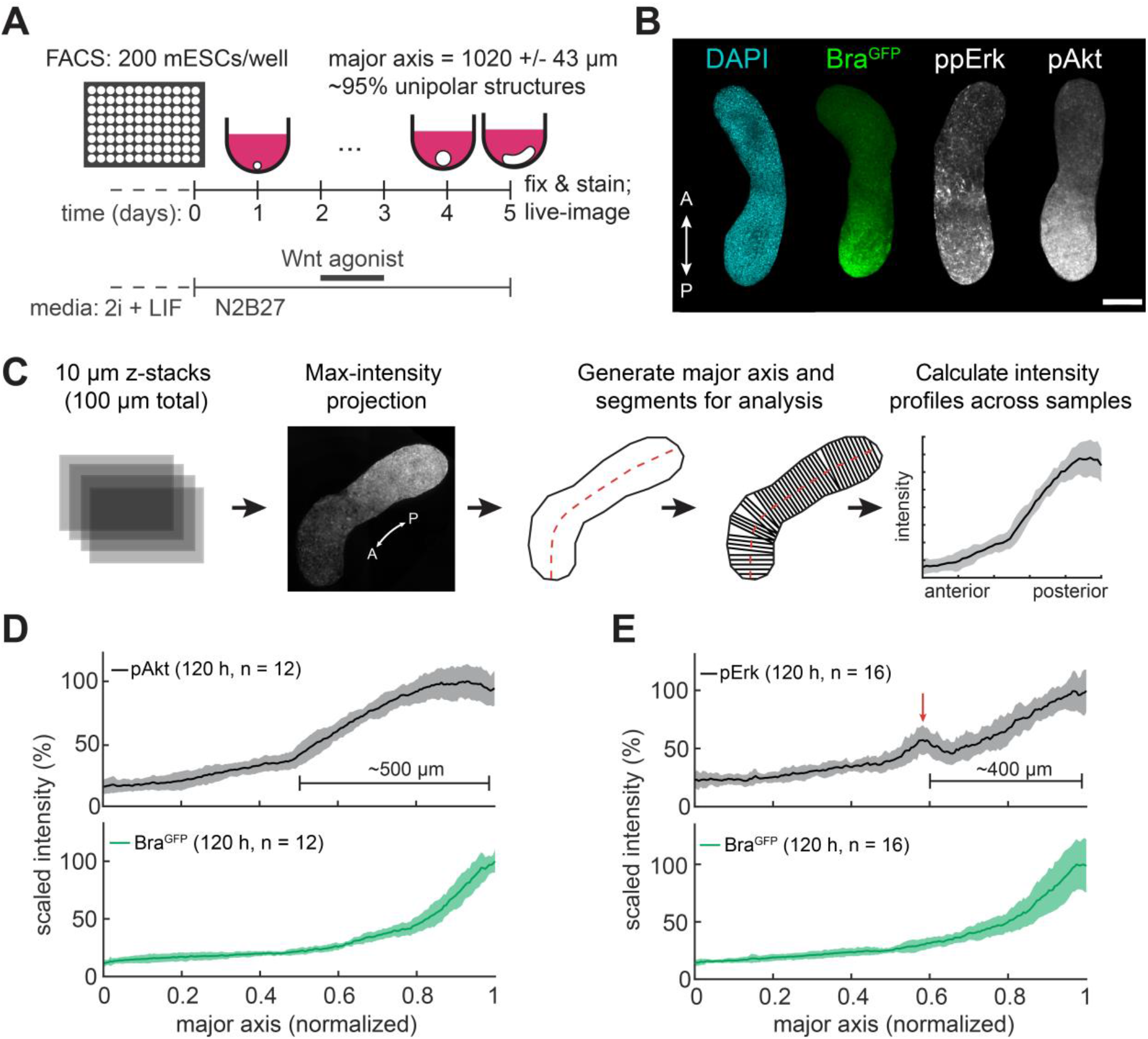
Spatial domains of Erk and Akt signaling during gastruloid elongation. (A) Schematic of gastruloid generation protocol. After seeding 200 cells per well, gastruloids are grown for 5 days, with addition of the Wnt agonist CHIR99021 from Days 2-3. By the end of Day 5, gastruloids elongate to ~1 mm in major axis length. (B) Representative gastruloid images showing nuclei (DAPI), Brachyury reporter expression (Bra^GFP^), and ppErk/pAkt signaling activity gradients. Scale bar is 200 μm. (C) Gastruloid imaging and quantification pipeline. Z-stacks of fixed gastruloids are maximum intensity projected and automatically segmented along their major axis for quantification of signaling patterns. (D-E) Spatial profiles of pAkt or ppErk immunofluorescence staining and Bra^GFP^ expression. The Erk secondary peak is indicated by the red arrow. For each analysis, 20 gastruloids were pooled, stained, and imaged. A number less than 20 is indicative of sample loss during the staining process. (N = 2 biological replicates). Shaded error regions represent standard deviation throughout the paper. For normalized major axes, 0 = anterior and 1 = posterior.

We used an image processing pipeline to quantify signaling patterns across multiple gastruloids (**Figure 1C; Methods**). In brief, 100 μm confocal image stacks were collected for each gastruloid, and the mean pixel intensity was obtained for each axial position, generating an intensity profile for each gastruloid along its major axis (**Figure 1C**). This quantification revealed long-range gradients in both Erk and Akt activity (**Figure 1D-E**) that declined steeply from the posterior domain to approximately the midpoint along the major axis (~500 μm absolute distance) and exhibited a shallower decline in the anterior half of the gastruloid. Erk and Akt activity were distributed over a broader range than Bra expression (**Figure 1D-E**). In addition, we found that the ppERK profile was non-monotonic, with a secondary peak in signaling activity located approximately 400 μm from the posterior pole (**Figure 1E**; red arrow). This is reminiscent of the ppErk intensity profile observed in the mouse embryo, where an oscillatory band of Erk activity marks the posterior-most somite^14^. Analysis of ppErk profiles for individual gastruloids (**Methods**; **Figure S1B**) reveals secondary ppErk peaks in 12 of the 16 samples analyzed (**Figure S1C**). The source of this variability could be explained by oscillatory Erk signaling, as observed during mouse somitogenesis, rather than a steady-state, fixed domain of activity.

### Kinase inhibitor treatments reveal functional roles for Erk/Akt signaling

How might gradients of Erk and Akt activity depend on upstream inputs, influence one another, and impact gastruloid morphogenesis? To begin to address these questions, we treated gastruloids with small-molecule inhibitors targeting the Erk pathway (the MEK inhibitor PD0325901), the Akt pathway (the PI3K inhibitor LY294002 and Akt inhibitor MK-2206), or FGFR itself (the FGFR inhibitor PD173074) and monitored their effects on signaling patterns and overall gastruloid morphogenesis (**Figure 2A**). Kinase inhibitors were added between days 4 and 5 of the protocol to examine their impact on axial elongation without disrupting earlier events such as the establishment of an A-P axis.

**Figure 2:**
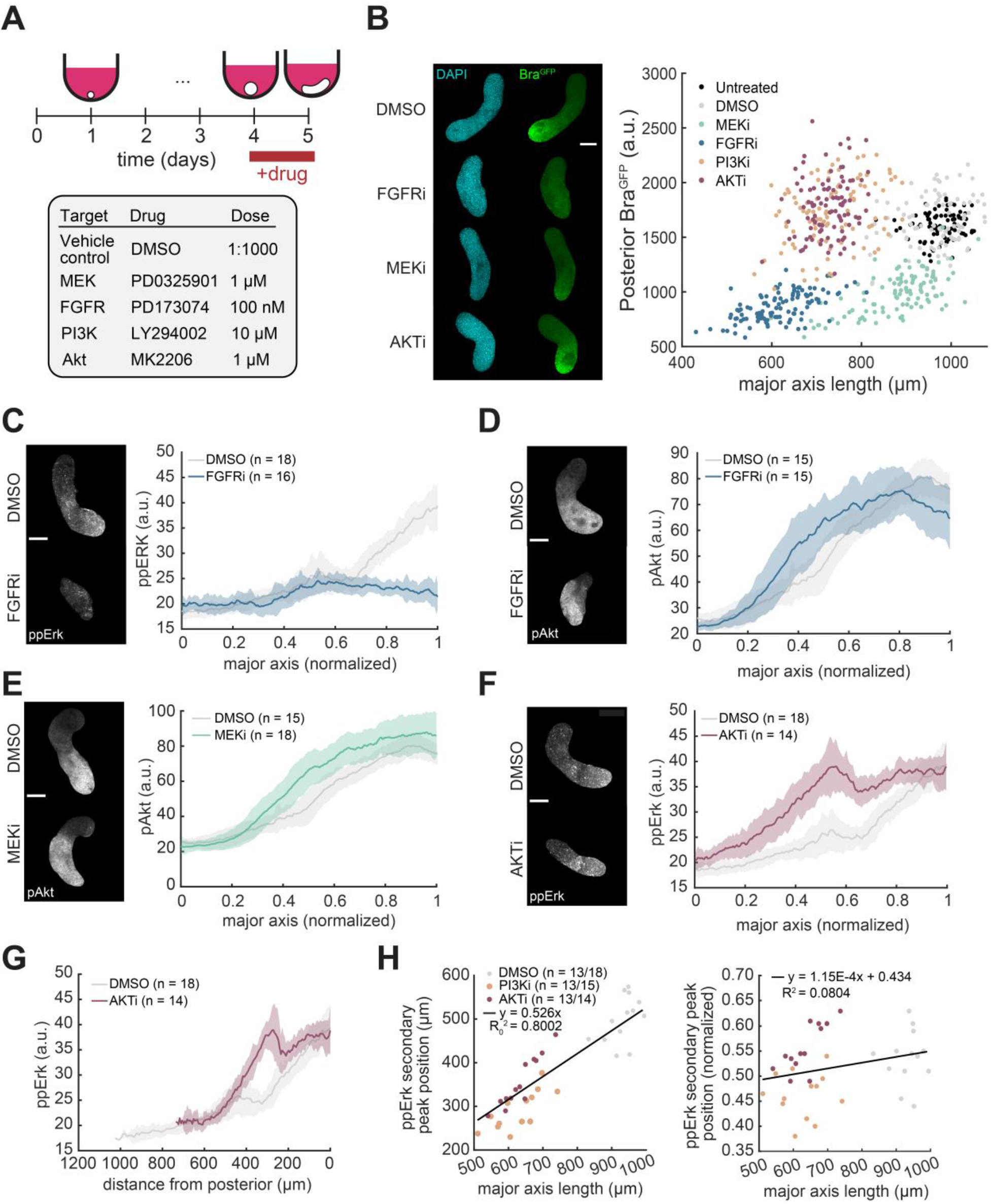
FGFR, Erk, and Akt signaling are required for proper gastruloid elongation. (A) Schematic of inhibitor treatment experiments. Inhibitors were administered during elongation from Day 4-5. Working concentration for each inhibitor is indicated. (B) Left panel: representative DAPI and Bra^GFP^ images for DMSO, FGFRi, MEKi, and AKTi treatments. Right panel: quantification of major axis length and Bra^GFP^ expression in the posterior 20% of the gastruloid after 24 h of inhibitor treatment, pooled from N = 2 biological replicates. Untreated n = 90. DMSO n = 92. MEKi n = 103. FGFRi n = 115. PI3Ki n = 101. AKTi n = 100. 120 gastruloids were seeded for each treatment condition, with a fraction lost during experimental manipulations. (C-F) Plots comparing either ppErk or pAkt spatial profiles between DMSO control and inhibitor-treated gastruloids, plotted on a normalized major axis where 0 is the anterior pole and 1 is the posterior pole. Each analysis began with 20 gastruloids. (N = 2 biological replicates). (G) Identical data to panel (F), but with ppErk intensity plotted as a function of absolute distance from the posterior pole. Note that the ppErk spatial profiles collapse onto the same curve with the exception of the secondary peak. (H) Left panel: distance of the secondary ppErk peak from the anterior pole as a function of major axis length for all individual DMSO and inhibitor-treated gastruloids with a detectable secondary Erk peak. The y-intercept for the linear regression is fixed at zero. Right panel: normalized position of the ppErk secondary peak as a function of major axis length. The 95% confidence interval for the slope of the linear regression is [-1.47×10^-5^, 2.46×10^-4^]. All scale bars are 200 μm.

FGFR inhibition resulted in the most profound defects, with a ~40% reduction in major axis length and complete absence of Bra^GFP^ enrichment in the posterior domain, consistent with the loss of Bra expression in the primitive streak of *Fgfr1*-/-embryos^23^ (**Figure 2B**). A similar loss in Bra^GFP^ expression was observed in response to MEK inhibitor treatment, with a partial reduction in gastruloid length that is consistent with prior reports^11,17,18,24^ (**Figure 2B**). In contrast, PI3K and Akt inhibition resulted in smaller gastruloids with a shorter major axis, yet had no impact on the posterior expression of Bra^GFP^ (**Figure 2B**). Akt inhibition closely phenocopied PI3K inhibition, demonstrating that Akt is the major pathway node downstream of PI3K in this system. Taken together, these results suggest that gradients of Erk and Akt activity play distinct roles in gastruloid elongation, with Erk required for both localized Bra expression and elongation, and Akt required for normal elongation but not mesodermal patterning.

Canonically, FGF stimulation is thought to activate both the Erk and Akt pathways, which may then cross-regulate one another *via* intracellular crosstalk^25^. To test whether similar principles apply in gastruloid elongation, we next measured Erk and Akt signaling gradients in each inhibitor condition. We first verified that MEKi and AKTi treatment indeed eliminated their respective Erk and Akt signaling gradients (**Figure S2C**; **Figure S2E**), validating that the inhibitors were functional at the concentrations used, whereas PI3Ki treatment reduced but did not eliminate Akt phosphorylation **(Figure S2F**). We next set out to test whether FGFR was required for both the Erk and Akt spatial patterns. Notably, while FGFRi treatment abolished the Erk gradient (**Figure 2C**; **Figure S2D**), we found that the amplitude and spatial range of Akt phosphorylation was unaffected (**Figure 2D**; **Figure S2H**). This result demonstrates that FGFR is required for the posterior-to-anterior gradient of Erk but not Akt. Control of Akt by other receptor tyrosine kinases (e.g. PDGFR) is well documented in other model systems^19,26^, and similar principles may also operate in the elongating gastruloid.

We tested for potential cross-talk between pathways by quantifying Erk patterning in Akt-inhibited gastruloids and *vice versa*. Just as in the case of FGFRi treatment, the Akt phosphorylation gradient was largely unaffected by the presence of a MEK inhibitor (**Figure 2E**; **Figure S2G**). The converse experiment – measuring Erk phosphorylation in the presence of an Akt or PI3K inhibitor – revealed an Erk pattern that was elevated relative to the DMSO-treated control at all normalized positions along the major axis (**Figure 2F**; **Figure S2A-B**). However, plotting the level of Erk activity as a function of *absolute* position along the major axis revealed similar overall gradients (**Figure 2G**), indicating that the primary effect of PI3K/Akt inhibition is to foreshorten the major axis, not alter the shape of the posterior-to-anterior ppErk gradient, with one exception noted below.

A close examination of ppErk staining across DMSO, AKTi and PI3Ki treatment conditions revealed that the secondary peak of Erk activity was always found at ~55% gastruloid length across all three treatment conditions, despite a roughly 25% reduction in major axis length between control and inhibitor-treated cases (**Figure 2F, Figure S2A-B**). These data suggested that the position of the secondary Erk peak might scale in proportion to overall gastruloid length. Indeed, plotting the ppErk signal as a function of absolute distance from the posterior pole demonstrated that the secondary peak is nearer to the posterior pole in the AKTi and PI3Ki treatment conditions (**Figure 2G, Figure S2A-B**). Unbiased measurement of the position of the ppErk peak in individual gastruloids across all three treatment conditions (**Methods**) revealed a linear scaling relationship between gastruloid length and the position of the Erk peak across and within treatment conditions (**Figure 2H**, left plot). No residual trend was observed when the normalized position of the Erk band was plotted against major axis length (**Figure 2H**, right plot). Thus, although gastruloids harbor non-interacting posterior-to-anterior gradients in Erk and Akt activity, superimposed upon this profile is a secondary peak in Erk activity near the midpoint of the gastruloid whose position robustly scales with overall gastruloid length.

In sum, quantification of the Erk and Akt signaling gradients across inhibitor-treated gastruloids revealed distinct regulatory and functional relationships for both signaling patterns. First, we find that both Erk and Akt activity are necessary for normal axis elongation, with inhibition of either signal reducing gastruloid length and/or Brachyury expression. Second, our experiments reveal that at least two upstream inputs define posterior-to-anterior signaling gradients: FGFR establishes the Erk activity pattern, whereas an unknown FGFR-independent input defines the Akt gradient. And third, we demonstrate that while the Erk and Akt gradients are largely non-interacting, at least one feature of the Erk pattern – the position of an activity peak near the gastruloid midpoint – scales with overall gastruloid length.

### Cell proliferation and cell-cell adhesion are spatially patterned in elongating gastruloids

How might patterns of Erk and Akt activity regulate gastruloid elongation? To begin to address this question, we first set out to define the cellular processes that drive unidirectional elongation in this system. We performed differential interference contrast (DIC) imaging on live gastruloids that were embedded in 50% Matrigel / 50% N2B27 to provide mechanical stability during imaging^10^ (**Figure 3A**; **Methods**). At the onset of elongation (96 h post-seeding), we observed the extrusion of a rigid cap-like domain (**Figure 3B,** left image; **Movie S1**); endpoint imaging confirmed that only this domain expressed Bra^GFP^, marking it as the nascent posterior domain. Particle image velocimetry (PIV) revealed that the posterior and anterior portions of the tissue flowed apart at a rate of ~10-15 μm/hour (**Figure 3B**, right image; **Methods**). Later, between 109-120 hours, the gastruloid exhibited steady, unidirectional elongation of the posterior domain (**Figure 3C,** left image; **Movie S2**). With the velocity of bulk tissue movement subtracted, PIV analysis demonstrated that cells delaminated from the pole and moved anteriorly at a rate of ~10-15 μm/h to drive overall outward displacement of the posterior cap (**Figure 3C**, right image). Kymographs of the elongating gastruloid revealed that the ~250 μm cap was extruded from the stalk at a rate of ~13 μm/h (**Figure 3D**), corroborating the results from the PIV analysis. These live-imaging data point to two processes underlying posterior elongation. First, the elongating posterior consists of a rigid cap and more fluid-like stalk, suggesting that differences in cell-cell adhesion and mechanical properties between these two tissue regions may play an important role. Second, posterior elongation occurs without a significant narrowing of the gastruloid minor axis, indicating that local cell proliferation must balance extrusion of the posterior domain to avoid a loss in tissue width.

**Figure 3:**
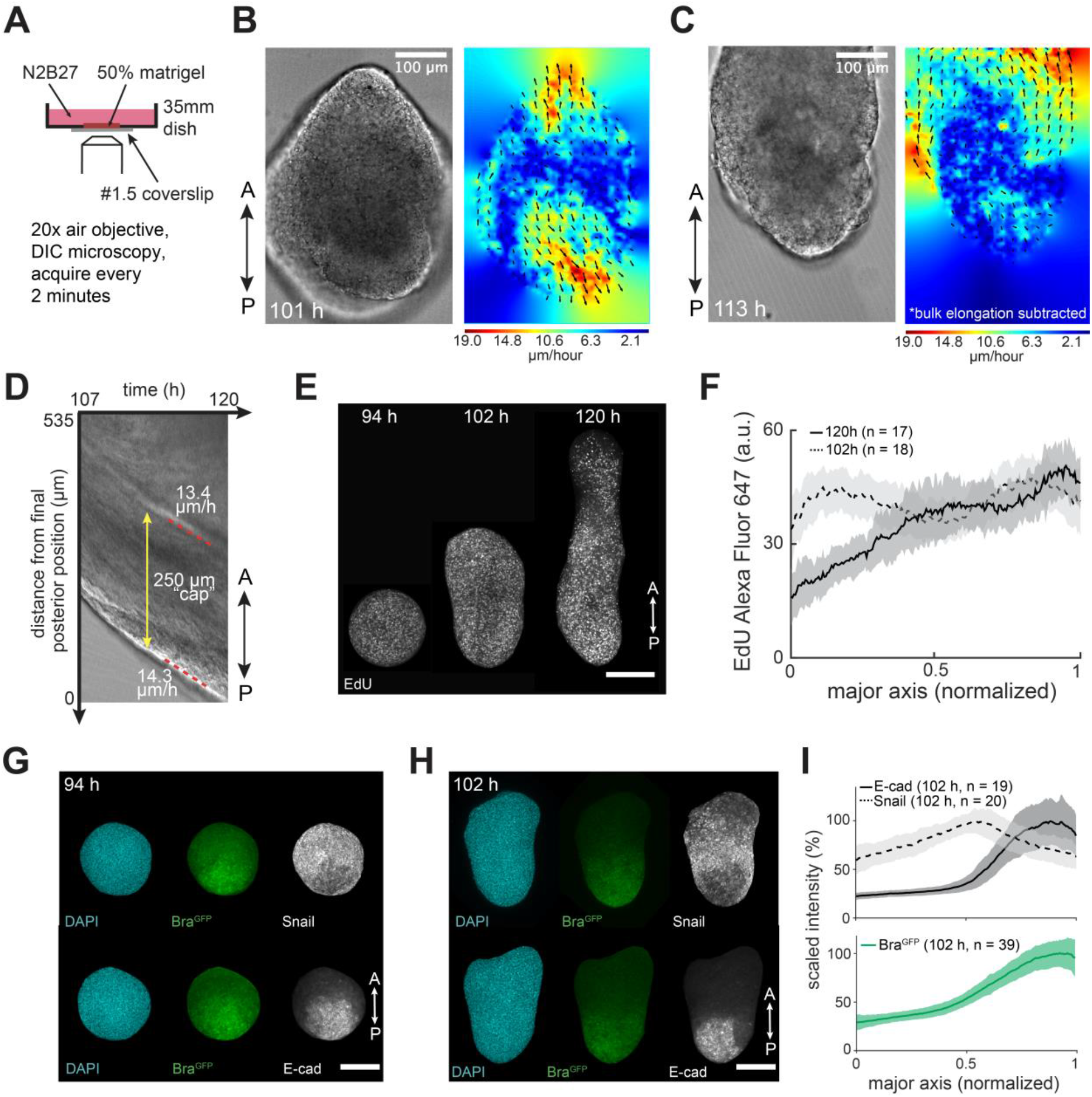
Characterizing mechanisms of gastruloid elongation. (A) Schematic of gastruloid live imaging protocol. Gastruloids are placed in 50% Matrigel to limit movement during live imaging. (B) Left panel: frame from Movie S1 at 101 hours after aggregation, in which the gastruloid has begun to extrude a posterior cap. (B) Right panel: mean velocity field generated from the previous hour of imaging using particle image velocimetry (PIV). Heat map represents magnitude of velocity vectors, with arrows indicating direction. (C) Left panel: frame from Movie 2 at 113 hours after aggregation. Right panel: PIV analysis from the previous hour of the timelapse, with the outward movement of the posterior tip subtracted. (D) Kymograph generated from a vertical line along the gastruloid major axis using the sample shown in (C). (E) Representative images of EdU staining at various timepoints during gastruloid elongation. (F) Quantification of EdU profiles for 102 h and 120 h gastruloids. Each analysis began with 20 gastruloids (N = 1). (G-H) Sample images of DAPI, Bra^GFP^, Snail, and E-cad at 94 h before physical elongation has begun (panel G), or 102 h, at the start of elongation (panel H). (I) Quantification of Bra^GFP^, Snail and E-cad spatial profiles at 102 h. The Snail and E-cad analysis began with 20 gastruloids each (N = 1). The Bra^GFP^ curve is aggregated for both samples. Scale bars are 200 μm unless labelled otherwise.

Can local differences in cell proliferation and cell-cell adhesion be directly observed? We first used an EdU staining assay to characterize local cell proliferation in elongating gastruloids (see **Methods**). Indeed, we found that while EdU staining was initially homogenous in unpolarized and early-elongating gastruloids, a posterior-to-anterior gradient in proliferation emerged as posterior elongation was established (**Figure 3E-F**). To characterize local differences in cell-cell adhesion, we stained for the canonical adhesion protein E-cadherin (E-cad) and the EMT-associated transcription factor Snail, factors whose expression often correlate with a tissue’s material properties due to their impacts on cell movement and cell-cell contact strength^27^. Consistent with prior studies^28,29^, we found that E-cad expression is restricted to the presumptive posterior just prior to gastruloid elongation, with Snail+ cells surrounding this domain (**Figure 3G**). Later during elongation, E-cad is expressed only at the posterior cap, whereas Snail protein forms a band just anterior to this cap (**Figure 3H-I**). Overall, our data support a model where posterior elongation is driven by two processes at the posterior cap: (1) localized cell proliferation at the posterior to support elongation without tissue thinning, and (2) continued posterior displacement of a rigid E-cad+ cap by newly added Snail+ cells.

### Perturbing Akt signaling alters cell proliferation and gastruloid size

We next sought to characterize how the gradient of Akt signaling might modulate either proliferation, cell-cell adhesion, or both. We began by measuring E-cad/Snail protein levels in elongating gastruloids treated for 24 h with the Akt inhibitor MK-2206 (AKTi) or vehicle control. We found that AKTi treatment resulted in smaller gastruloids but had little impact on the spatial patterns of Snail (**Figure 4A**) or E-cad expression (**Figure S3A**). Notably, we again found evidence of pattern scaling with overall gastruloid size: the Snail peak was observed at ~60% along the major axis in both control and AKTi conditions (**Figure 4A**), and plotting the position of peak Snail expression as a function of major axis length for individual control and inhibited gastruloids revealed a linear relationship (**Figure 4B**). FGF/Erk signaling is known to regulate Snail expression^23^, suggesting that size-invariant scaling of the secondary Erk peak may also be important for patterning downstream effector genes such as Snail. More broadly, our observation that AKTi treatment did not alter the overall patterns of E-cad, Snail, or Bra^GFP^ expression (**Figure 2B**) further supports a model where Akt signaling is dispensable for much of the patterned gene expression along the gastruloid’s A-P axis.

**Figure 4:**
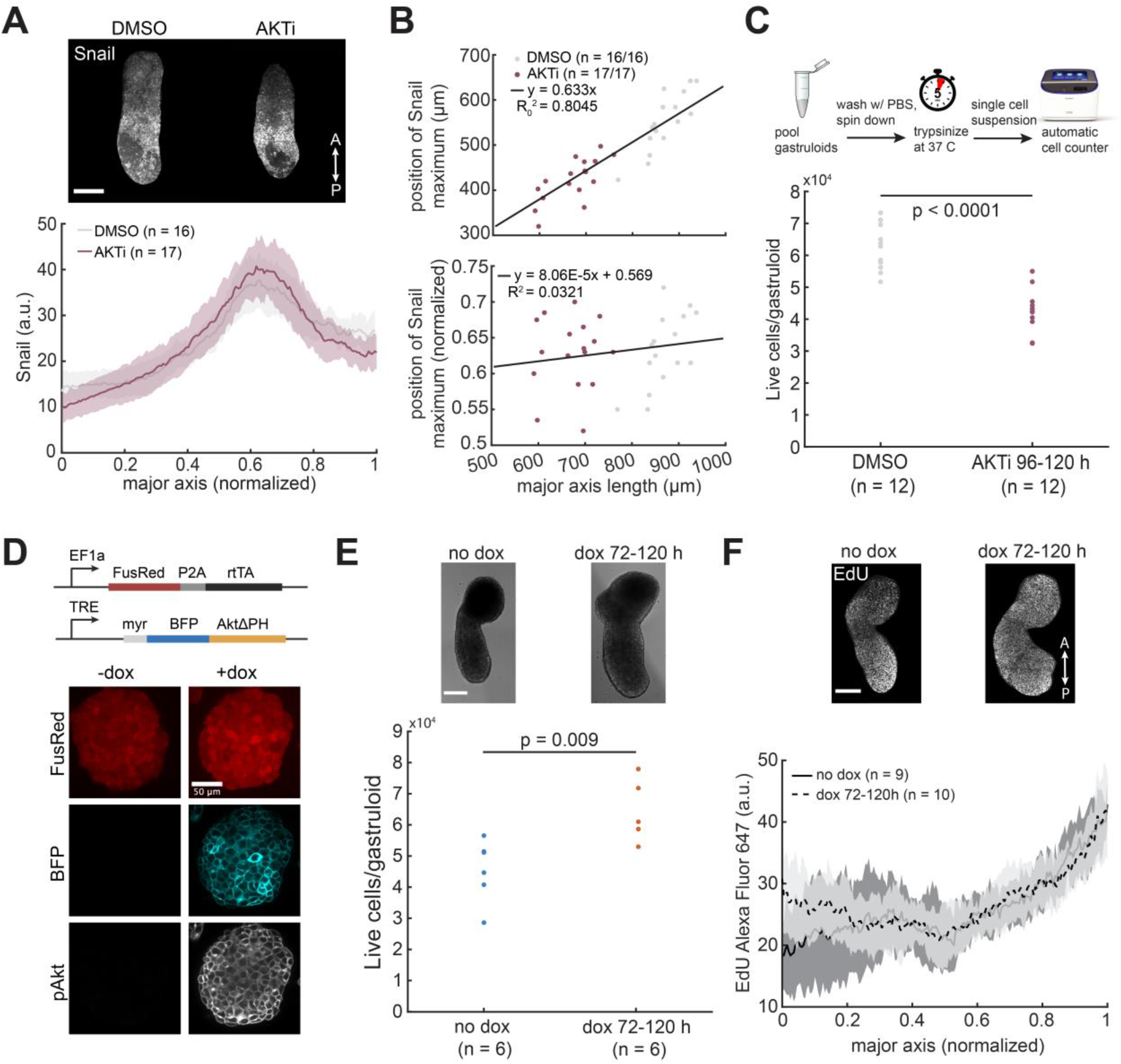
PI3K/Akt signaling modulates proliferation along the gastruloid major axis. (A) Spatial profile of Snail expression after DMSO or AKTi treatment. Analysis began with 20 gastruloids each (N = 2). (B) Top panel: distance of the Snail peak from the anterior pole as a function of major axis length for all DMSO and AKTi-treated gastruloids. The y-intercept for the linear regression is fixed at zero. Bottom panel: normalized position of the Snail peak as a function of major axis length. The 95% confidence interval for the slope of the linear regression is [-8.16×10^-5^, 2.43×10^-4^]. (C) Live cells per gastruloid for Akt inhibition compared to vehicle control. Each data point represents the average cell count per gastruloid found after pooling 5 gastruloids. Data is aggregated from N = 3 biological replicates. (D) Schematic of dox-Akt expression system and representative images for FusionRed, BFP, and pAkt in the absence or presence of 1 μg/mL doxycycline. (E) Live cells per gastruloid after treatment with 1 μg/mL doxycycline from 72-120 h, compared to a doxycycline-free control. Each data point represents the average cell count per gastruloid found from pooling 10 individual gastruloids. Data is aggregated from N = 3 biological replicates. (F) EdU staining after treatment with 1 μg/mL doxycycline from 72-120 h, compared to a doxycycline-free control. Each analysis began with 10 gastruloids (N = 3). All scale bars are 200 μm unless labelled otherwise.

Might Akt instead act to regulate cell proliferation? To test this possibility, we developed a protocol to pool and dissociate known numbers of gastruloids and measure both the total and live cell counts per gastruloid (**Methods**; **Figure 4C**, upper schematic). We found that AKTi treatment led to a ~30% reduction in live cells per gastruloid at 120 h, from an average of 62,000 cells per gastruloid in the vehicle control case to 43,000 cells per gastruloid in the AKTi case (**Figure 4C**, lower panel). The reduced cell count could in principle arise from either a decrease in cell proliferation or an increase in cell death, but trypan blue staining revealed only a minor increase in the proportion of dead cells after AKTi treatment (**Figure S3B-C**). Akt inhibition thus appears to decrease gastruloid size primarily by suppressing cell proliferation.

We next set out to perform the converse experiment: testing whether global Akt hyperactivation might induce hyper-proliferation and increased gastruloid growth. As an initial test of this model, we treated gastruloids with platelet derived growth factor (PDGF), a known activator of Akt signaling in a wide variety of developmental contexts^19,26^. We confirmed that, in gastruloids, PDGF treatment leads to a global increase in pAkt levels, most notably in the anterior domain where Akt activity is normally absent (**Figure S3D**). In conjunction with this expanded profile of Akt signaling, we observed high levels of EdU staining in the anterior domains of elongating gastruloids (**Figure S3E**), indicative of a shift from local to global cell proliferation, as well as significant gastruloid enlargement in response to PDGF treatment (**Figure S3F**).

Receptor-level activation using PDGF activates a number of downstream pathways, including PI3K/Akt, Ras/Erk, PLCγ, and JAK/STAT signaling^26^. We thus sought an alternative method to more specifically activate Akt. Inspired by chemogenetic and optogenetic tools demonstrating that Akt membrane localization is sufficient to drive its activation^30,31^, we engineered a stable mESC cell line with doxycycline-inducible expression of BFP-tagged, membrane-localized Akt, hereafter referred to as dox-Akt cells (**Figure 4D**). We confirmed that 24 h treatment with 1 μg/mL doxycycline drove a dramatic increase in both Akt phosphorylation and BFP expression in dox-Akt cells (**Figure 4D**), demonstrating that this cell line can indeed be used to robustly produce high levels of Akt activity in response to doxycycline treatment. We found that when dox-Akt gastruloids were exposed to 1 μg/mL of doxycycline from 72-120 h, the aggregates were wider and longer (**Figure S3G**) and contained more overall cells (**Figure 4E**) without a change in gastruloid aspect ratio (**Figure S3H**) or cell viability (**Figure S3I**). Akt hyperactivation also drove increased cell proliferation at the anterior domain of the gastruloid, as marked by EdU staining (**Figure 4F**). Finally, just as in the case of AKTi treatment, the domains of Snail and E-cad expression were unperturbed in dox-Akt gastruloids (**Figure S3J-K**). These experiments demonstrate that Akt signaling regulates cell proliferation in elongating gastruloids, and, taken together with our observation of a posterior-to-anterior Akt activity gradient, suggest that local Akt activity may serve to drive local cell production at the posterior and support the volumetric increase of this domain.

### Perturbing Erk activity alters gene expression patterns and gastruloid shape

Finally, we measured how Erk signaling affected cellular processes associated with gastruloid elongation. In contrast to our previous observations with Akt inhibition (**Figure 4A, Figure S3A**), treating gastruloids with the MEK inhibitor PD0325901 (MEKi) resulted in a dramatic change in the patterns of E-cad and Snail expression. Snail expression was dramatically reduced in MEKi-treated gastruloids (**Figure 5A**) and was replaced by an extended domain of E-cad expression to approximately ~50% along the gastruloid major axis (**Figure 5B**). MEKi-treated gastruloids also had a lower average number of cells than the vehicle control, but the effect was less pronounced than with Akt inhibition (**Figure S4**). Thus, unlike Akt, Erk activity directly impacts the patterned expression of genes associated with cell-cell adhesion during gastruloid elongation.

**Figure 5:**
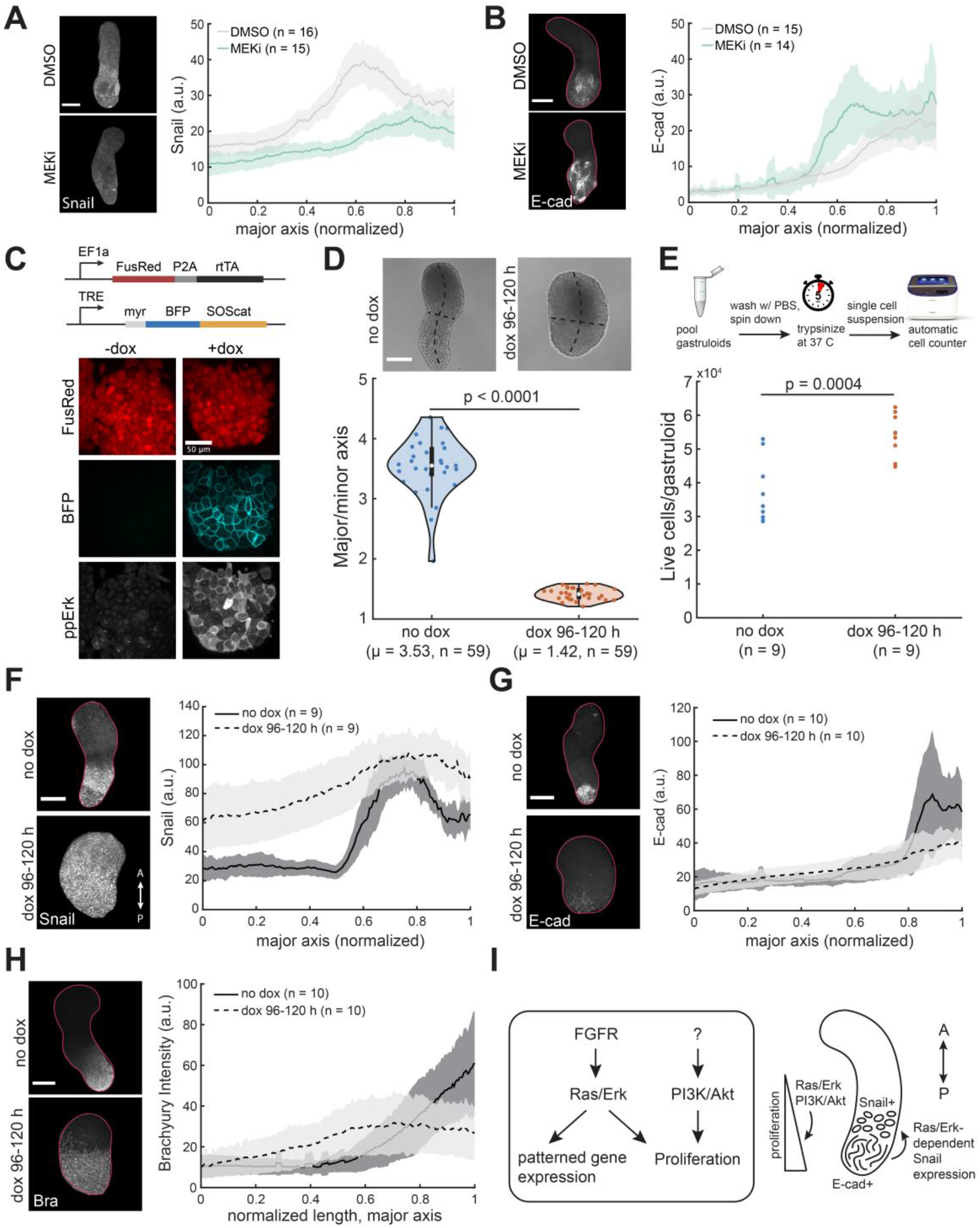
Ras/Erk signaling is necessary and sufficient for Snail expression. (A-B) Spatial profiles of Snail and E-cad expression after MEK inhibition, compared to vehicle control. Analysis began with 20 gastruloids each (N = 2). (C) Schematic of dox-SOScat expression system and representative images for FusionRed, BFP, and ppErk in the absence or presence of 1 μg/mL doxycycline. (D) Major/minor aspect ratio for dox-SOScat gastruloids treated with 1 μg/mL doxycycline from 96-120 h, compared to a doxycycline-free control. Each treatment began with 60 gastruloids. N = 3 biological replicates. (E) Live cells per gastruloid after treatment with 1 μg/mL doxycycline from 96-120 h, compared to a doxycycline-free control. Each data point represents the average cell count per gastruloid found from pooling 10 individual gastruloids. Data from N = 3 biological replicates. (F-H) Spatial profiles of Snail (panel F), E-cad (panel G), and Brachyury (panel H) for dox-SOScat gastruloids treated with 1 μg/mL doxycycline from 96-120 h treatment, compared to a doxycycline-free control. Analysis began with 10 gastruloids each (Snail and E-cad, N = 3; Brachyury, N = 1). (I) Schematic demonstrating the functionality of Ras/Erk and PI3K/Akt signaling gradients during gastruloid elongation. All scale bars are 200 μm unless labelled otherwise.

We performed time-lapse imaging of MEKi-treated gastruloids to better understand how altered domains of Snail and E-cad expression might affect elongation (**Movie S3**). The resulting movies revealed that, although elongation was still possible, the posterior cap became longer and more pointed over time (**Figure 5A-B**, lower images; **Movie S3**). Analysis of multiple time-lapse movies revealed a significant difference in the gastruloid tip’s radius of curvature after long-term MEKi treatment, with no difference at the beginning of the experiment just after Matrigel embedding (**Figure S5A-B**). Similarly, a comparison of kymographs revealed a gradual enlargement of the posterior cap and a reduction in the velocity of the cap’s posterior movement in MEKi-treated gastruloids (**Figure S5C**), consistent with the observation of an extended E-cad+ domain in these conditions (**Figure 5B**). Erk inhibition is thus likely to disrupt tissue dynamics at least in part by increasing the size of the E-cad+ posterior cap over time, leading to defects in its shape and outward movement, potentially due to increased cell-cell adhesion. Our results are also consistent with observations in the mouse embryo, where conditional inactivation of *Fgf4/8* during posterior elongation results in a decrease in *Snail* expression and a narrowing of the tail bud when compared to control embryos^13^.

We also engineered gastruloids in which Erk signaling could be globally activated on demand using a doxycycline-inducible, membrane-localized activator of Ras (dox-SOScat) (**Figure 5C**), analogous to the dox-Akt system described earlier. SOS is a guanine nucleotide exchange factor that, when recruited to the membrane, activates Ras, and localizing the catalytic domain of SOS to the membrane is sufficient to potently activate the Ras/Erk signaling cascade^32,33^. Treatment of clonal dox-SOScat mESCs with 1 μg/mL doxycycline for 24 h resulted in elevated BFP expression and ppErk levels, validating the efficacy of this approach (**Figure 5C**). Treating dox-SOScat gastruloids with doxycycline from 96-120 h resulted in a profound morphological change: the resulting gastruloids were nearly spherical, with a dramatically reduced major axis and expanded minor axis length (**Figure 5D**). Despite the reduction in major axis length, dox-SOScat gastruloids contained more cells (**Figure 5E**), confirming Ras/Erk signaling as a proliferative signal in this context.

We hypothesized that the near-isotropic growth of dox-SOScat gastruloids was due to a substantial loss of polarized gene expression, disrupting local control over cell mechanics. Indeed, staining for Snail and E-cad in dox-SOScat gastruloids treated with doxycycline from 96-120 h revealed global expression of Snail and a near-complete loss E-cad expression (**Figure 5F-G**). Ras/Erk hyperactivation during gastruloid elongation led to more complex changes to the pattern of Bra protein expression: we observed both an anterior expansion of the Brachyury-positive domain, as well as a decrease in maximum Bra protein levels at the posterior pole (**Figure 5H**). These data are consistent with data from the chick embryo in which overexpression of either FGF8 or constitutively-active MEK1 leads to an expansion but not global induction of Bra expression^15,34^. Taken together, our data indicates that spatially patterned Erk signaling plays a crucial role in producing localized domains of gene expression and tissue architecture in elongating gastruloids. Loss of Erk activity eliminates Bra and Snail expression and reduces posterior elongation, whereas Erk hyperactivation globally activates Snail expression and drives isotropic growth.

## Discussion

Mammalian gastrulation involves a multitude of processes that occur in tight coordination. Cells proliferate, undergo morphogenetic movements that expose them to multiple potentially time-varying extracellular cues, and execute fate decisions based on these cues. Embryonic stem cell-derived gastruloids provide an excellent reduced-complexity model to dissect the constituent processes that underlie mammalian posterior morphogenesis. They are highly amenable to genetic and chemical perturbations, can be generated in large numbers with reproducible outcomes, and exhibit three-dimensional signaling patterns, cell movements, and cell fate specification events, enabling the researcher to dissect how these processes might interact.

Posterior tailbud elongation, a central process in vertebrate development, is partially recapitulated in mESC-derived gastruloids as a striking morphological change on the fifth day of culture. This reduced-complexity model allows us to address many open questions around how signaling gradients along the anterior-posterior axis control axial elongation. We found that the kinases Erk and Akt constitute two such patterned signals, forming largely independent posterior-to-anterior activity patterns with similar spatial ranges that go on to regulate cell proliferation, patterned gene expression, or both. Overall, our study provides a quantitative map of two natural signaling gradients in an emerging model of mammalian gastrulation and details how disrupting these patterns alters tissue morphogenesis.

Careful quantification of Erk and Akt activity in the presence of kinase inhibitors revealed some unexpected relationships among these stimuli (**Figure 5I**). For example, we found that while FGFR inhibition eliminated the Erk pattern, it had no effect on the posterior-to-anterior gradient of Akt activity (**Figure 2**). This result suggests that Akt is normally activated by a distinct upstream ligand and receptor, possibly reflecting gradients of other receptor tyrosine kinase ligands (e.g. PDGF)^19,26^. In addition, we report an unexpected scaling relationship in one feature of the Erk signaling pattern with overall A-P axis length: a secondary peak of Erk activity that appears near the middle of the gastruloid. We show that the position of this secondary peak scales linearly with A-P axis length even after Akt or PI3K inhibition (**Figure 2**), a perturbation that shortens the A-P axis by roughly 25%. Similar scaling is also observed for the Erk target gene Snail (**Figure 4**), further supporting a model whereby Erk pattern scaling supports proper subdivision of the gastruloid’s posterior region into distinct domains of gene expression. In future work it would be interesting to further define the functional circuits that produce this pattern scaling relationship and to examine the consequences of its disruption.

Our data indicates that Erk and Akt exert somewhat separable effects on gastruloid morphogenesis. Experiments that either inhibited or hyperactivated Akt resulted in gastruloids with altered cell counts, suggesting that an Akt activity gradient titrates overall gastruloid size by tuning local proliferation rates. In addition to regulating cell proliferation, perturbations to Erk signaling profoundly altered the expression patterns of multiple genes: the mesoderm-associated transcription factor Brachyury, the EMT-associated transcription factor Snail, and the cell adhesion molecule E-cadherin. As might be expected for perturbations in genes involved in cell movement and adhesion, global perturbations to Erk activity altered gastruloid morphology, producing an elongated but pointed posterior domain when Erk is globally inhibited, and isotropic gastruloid growth when Erk is globally activated. These results might be summarized by a simple conceptual framework: both Akt and Erk signaling control gastruloid size, whereas Erk activity also regulates the shape of the growing tissue.

Nevertheless, many aspects of signaling regulation over gene expression patterns are still mysterious. For example, whereas global, ectopic Erk activation produces global Snail expression, in unperturbed gastruloids Snail is only present at a subset of locations where Erk activity is high (e.g., the central stripe, but not the posterior tip). This paradox might be resolved by a careful study of the dynamics of Erk activation, Snail expression and cell movement – for example, Erk activity might precede both anterior cell migration and Snail expression, so cells that are initially Erk-high at the posterior pole only express Snail once they have moved anteriorly. Additionally, we find that the relationship between Erk activity and Brachyury expression is complex: in response to global Erk activation, Bra expression increases in the middle of the gastruloid but decreases at the posterior tip. FGFR/Erk signaling is only one of many patterned stimuli in an elongating gastruloid, and it is likely that some of this complexity is provided by interactions with additional signals that are also patterned along the anterior-posterior axis such as Wnt and Nodal signaling^9^. Finally, dynamic Erk activity has been noted in an increasing number of model systems, including pulses of Erk activity that correlate to cell fate acquisition in the early mouse embryo^35,36^. The observation of salt-and-pepper Erk staining in elongating gastruloids may also be explained by asynchronous pulses, suggesting that much may be revealed by a careful study of the prevalence and function of signaling dynamics in gastruloid development.

A quantitative atlas of developmental signaling patterns can serve many useful functions. We have shown that in conjunction with perturbations, quantifying signaling patterns can reveal regulatory and scaling relationships between the patterns. Such quantification is also a prerequisite to “rewriting” patterns using the tools of optogenetics or synthetic biology. We must have an accurate understanding of where and when a particular signaling pattern is active, and what role it plays, before we can predict how an alternative pattern might drive novel outcomes. Many optogenetic tools are already available for controlling Erk, Akt and receptor tyrosine kinase activity in cells and organisms^31,33,37,38^, and we anticipate that the data presented here may enable the experimentalist to explore the rules of morphogenesis through delivery of alternative patterning cues.

## Methods

### Plasmids and cloning

All constructs were cloned into a PiggyBac plasmid (System Biosciences). Linear DNA fragments were amplified via PCR using CloneAmp HiFi PCR premix (Takara Bio, 639298). PCR products were cut from agarose gels and purified using the Nucleospin gel purification kit (Takara Bio, 240609). Final plasmids were constructed using In-Fusion HD (Takara Bio, 638910) and amplified in Stellar chemically competent E. coli (Takara Bio, 636763). DNA was extracted by miniprep (Qiagen, 27104). All plasmid verification was performed by Sanger sequencing (Genewiz) or nanopore sequencing (Plasmidsaurus). The Akt gene (without the pleckstrin homology domain) was codon optimized for Mus musculus and was ordered as a gBlock (Integrated DNA Technologies). The SOScat gene was cloned from existing plasmids and can be obtained on Addgene (Plasmid #86439). The mSnail gene was ordered from Addgene (Plasmid #34583).

### Routine cell culture

The E14TG2a Bra^GFP^ cell line was a gift from Dr. Alfonso Martinez Arias (Cambridge University). The dox-inducible myr-Akt and dox-inducible myr-SOScat cell lines were generated using wild-type E14TG2a cells (ATCC). All other experiments were carried out using the E14TG2a Bra^GFP^ cell line.

All mESCs were grown in 2i + LIF media. This consists of a base media of GMEM (Millipore Sigma, G6148) with 10% ESC qualified fetal bovine serum (R&D Systems, S10250), 1:100 GlutaMAX (Gibco, 35050-061), 1:100 MEM non-essential amino acids (Gibco, 11140-050), 1 mM sodium pyruvate (Gibco, 11360-070), and 100 μM 2-mercaptoethanol (Gibco, 21985-023). For a working stock of 2i + LIF media, this base media is supplemented with 1:100 Pen Strep (Gibco, 15140-122), 10^3^ units/mL ESGRO recombinant mouse LIF protein (Millipore Sigma, ESG1107), 2 μM PD0325901 (Tocris, 4192), and 3 μM CHIR99021 (Tocris, 4423).

mESCs were grown in filter-capped tissue culture flasks. These flasks were treated with a 0.1% gelatin solution in water (Millipore Sigma, ES-006-B) for 30 minutes prior to plating cells. Cells were passaged every 48 hours. To passage, media was aspirated from the flask and cells were washed with pre-warmed DPBS (-)Calcium (-)Magnesium (Gibco, 14190144). The DPBS was aspirated and cells were treated with pre-warmed TrypLE Express Enzyme, enough to evenly cover the bottom of the flask (Gibco, 12605028). Cells were incubated in trypsin at 37°C for 5 minutes, then 2i + LIF media was added and cells were triterated to form a single-cell suspension. Cells were centrifuged at 135 relative centrifugal force for 5 minutes, then the supernatant was aspirated and the cell pellet was resuspended in 2i + LIF media. The 0.1% gelatin solution was aspirated from the flask, and cells were plated at the desired density (typically a 1:5 to 1:10 dilution). Cell lines were routinely tested for mycoplasma using a PCR-based universal mycoplasma detection kit (ATCC, 30-1012K).

### Cell line generation

Clonal transgenic cell lines were generated via transfection and PiggyBac genomic integration. Cells were plated 24 hours before transfection in a cell-culture treated 6-well plate (Fisher Scientific, 14-832-11). To transfect, 2000 ng of the PiggyBac vector plasmid and 500 ng of the PiggyBac transposase plasmid (System Biosciences, PB210PA-1) were mixed in 125 μL Opti-MEM (Gibco, 31985-070). 5uL of lipofectamine stem transfection reagent (Invitrogen, STEM00001) was mixed in 125 μL Opti-MEM. The diluted DNA and transfection reagent were then combined, mixed thoroughly, and allowed to incubate at room temperature for 30 minutes before gently pipetting onto the cells. Cells were not passaged for at least 24 hours after transfection.

4 days after transfection, cells were run on a Sony SH800 cell sorter (Sony Biotechnology). Single cells positive for the fluorescent selection marker were sorted into a flat-bottom plastic 96-well plate (Corning, 353072) for the generation of clonal lines. These plates were left in the incubator for 7 days before screening colonies for fluorescent expression and morphology. Once propagated to a sufficient cell number, clonal lines were assayed for transgene functionality and for their ability to grow healthy gastruloids.

### Gastruloid protocol

Gastruloids were grown in N2B27 media. This consists of a 1:1 mixture of DMEM/F-12 (Gibco, 11320033) and neurobasal medium (Gibco, 21103049), supplemented with 100 μM 2-mercaptoethanol, 1:100 N-2 (Gibco, 17502048), 1:50 B-27 (Gibco, 17504044), and 1:100 pen strep. Media was sterile-filtered using 0.22 μm pore size Steriflip 50 mL conicals (Millipore Sigma, SCGP00525).

Gastruloid formation followed a standard protocol^39^, one variation being that here we used a cell sorter to seed gastruloids as opposed to a hemocytometer / multichannel pipette. In short, a single-cell suspension of mESCs was washed twice in DPBS (-)Calcium (-)Magnesium to remove any trace amounts of 2i + LIF media. This washed pellet was then resuspended in prewarmed N2B27 media and run of a Sony SH800 cell sorter. The events were gated to exclude doublets and cell debris. Cells were sorted into a prepared 96-well round-bottom ultra-low attachment microplate (Corning, 7007). To minimize the effects of evaporation at the edges of the plate, the perimeter wells were filled with 100 μL water containing 1:100 pen strep. The inner 60 wells were then filled with 40 μL N2B27. 200 single mESCs were sorted into each of these 60 inner wells. The plate was then immediately returned to the incubator.

After 48 hours, 150 μL pre-warmed N2B27 with 3 μM CHIR99021 is added to each gastruloid-containing well. At the 72-hour mark, 150 μL of media is removed from each well and replaced with 150 μL fresh N2B27. Likewise, at the 96-hour mark, 150 μL of media is removed from each well and replaced with 150 μL fresh N2B27. All inhibitor treatments were carried out between 96-120 hours after aggregation. The inhibitors used in this study are as follows: PD173074 (Selleck Chem, S1264), PD0325901 (Tocris, 4192), LY294002 (Cell Signaling Technology, 9901), and MK-2206 (Selleck Chem, S1078). For global PI3K/Akt activation, platelet-derived growth factor-BB (Millipore Sigma, P3201) was used at 100 ng/mL. All doxycycline (Millipore Sigma, D9891) treatments were at a final concentration of 1 μg/mL.

### Immunofluorescence staining

To stain adherent mESCs, media was aspirated and 4% paraformaldehyde in PBS (Fisher Scientific, AAJ19943K2) was added to fix the cells. After 15 minutes at room temperature, cells were washed with DPBS (-)Calcium (-)Magnesium, 3 times for 5 minutes per wash. DPBS was aspirated and PBSFT was added for one hour at room temperature. PBSFT consists of 10% fetal bovine serum and 0.2% Triton X-100 (Millipore Sigma, 648466) in DPBS. After one hour, blocking buffer was aspirated and the primary antibody, diluted in PBSFT, was added to the cells. Cells were incubated in the primary antibody solution overnight at 4 °C. The next day, the primary antibody solution was aspirated and the cells were washed 3 times in DPBS, 5 minutes per wash. The secondary antibody, diluted in PBSFT, was added to the cells and let incubate for 90 minutes at room temperature. Cells were then washed another 3 times with PBS before imaging.

Staining gastruloids follows a similar procedure^39^. 150 μL of media was removed from each well and replaced with 150 μL of 4% PFA in PBS. Gastruloids were fixed at 4°C for 2 hours, rocking continuously. 150 μL was then removed from each well and replaced with 150 μL of DPBS (-)Calcium (-)Magnesium. This wash step was repeated for a total of 3 washes. Following the washes, gastruloids that are undergoing the same primary/secondary antibody staining regiment are pooled into a single well of the round-bottom 96-well plate. Here, a 200 μL pipette with the tip cut off is used to transfer gastruloids under a dissection microscope. Once gastruloids are pooled (typically 10 gastruloids per well), the PBS is completely removed from the well and replaced with PBSFT. Gastruloids are incubated in PBSFT overnight at 4°C. The following day, the PBSFT is removed from each well and replaced with the primary antibody diluted in PBSFT. Gastruloids are incubated in the primary antibody overnight at 4°C. The following day the primary antibody is washed out using PBSFT. This consists of 3 short washes (~5 min intervals between washes) followed by 3 long washes (~1 h). After the final wash, the PBSFT is completely removed and replaced with the appropriate secondary antibody diluted in PBSFT. This is incubated overnight, and the following day the same wash procedure (3 short, 3 long washes) is carried out. For any samples that were stained with DAPI (Thermo Fisher, 62248), 1 μg/mL DAPI was added with the secondary antibody.

Primary antibodies used in this work: phospho-p44/42 MAPK, Erk1/2 phosphorylated at Thr202 and Tyr204 rabbit (Cell Signaling Technology, 4370, 1:200 dilution), phospho-Akt at Ser473 rabbit (Cell Signaling Technology, 9743, 1:400 dilution), Snail C15D3 rabbit (Cell Signaling Technology, 3879, 1:50 dilution), E-cadherin 24E10 rabbit (Cell Signaling Technology, 2606, 1:200 dilution), and Brachyury goat (R&D Systems, AF2085, 10 μg/mL working concentration). Secondary antibodies: Goat anti-rabbit Alexa Fluor 647 conjugate (Invitrogen, A27040, 1:500 dilution), chicken anti-goat Alexa Fluor 647 conjugate (Invitrogen, A21469, 1:500 dilution), and goat anti-rabbit Alexa Fluor 405 conjugate (Invitrogen, A31556, 1:500).

### EdU Staining

5-ethynyl-2’-deoxyuridine (EdU) staining was performed using the 647 EdU Click Proliferation Kit (BD Biosciences, 565456). First, a 40 μM solution of EdU was made in prewarmed N2B27. 95 μL of N2B27 was removed from each gastruloid-containing well, and 95 μL of N2B27 containing EdU was added for a final concentration of 20μM EdU in the well. The plate was returned to the incubator for 2 hours to allow time for the EdU to integrate into S phase cells. After 2 hours, 150 μL of media was removed from each well and 150μL of DPBS (-)Calcium (-)Magnesium was added to wash. 150 μL was then removed from each well and replaced with 150 μL of 4% paraformaldehyde in PBS. Fix the gastruloids, rocking at 4°C, for 1 hour. After fixation, wash 3 times with DPBS (-)Calcium (-)Magnesium. Then, as described previously, pool gastruloids of the same treatment conditions into one well of the 96 well plate (typically 10 gastruloids/well). Once gastruloids are pooled, remove all liquid and replace with saponin-based permeabilization buffer. Incubate overnight at 4°C. The following day, make a working solution of the red 645 azide cocktail. For a final volume of 500 μL, first dilute 1 μL of the azide dye into 24 μL DMSO. Into 435 μL DPBS (-)Calcium (-)Magnesium, add 10 μL of the copper catalyst solution, then 5 μL of the diluted dye azide, then 50 μL of the buffer additive. Taking the plate of gastruloids, remove all PBSFT from the gastruloid-containing wells. Add 100 μL of the azide cocktail to each well. Pipette up and down in the well to ensure thorough mixing of the cocktail around the gastruloids. Incubate for 1 hour, rotating at room temperature and protected from light, mixing the solution again after 30 minutes. Once the incubation is complete, remove the azide cocktail and wash 3 times with the saponin-based permeabilization buffer.

### Microscopy and staining quantification

All images in this work were acquired on a Nikon Eclipse Ti confocal microscope with a Prior linear motorized stage, a Yokogawa CSU-X1 spinning disk, an Agilent laser line module containing 405, 488, 561 and 650nm lasers, and an iXon DU897 EMCCD camera.

For adherent cells, such as the dox-inducible myr-Akt and myr-SOScat lines, cells were plated on a 96-well glass bottom plate with high performance #1.5 cover glass (Cellvis, P96-1.5H-N) and were imaged using a 40x oil emersion objective. To image fixed gastruloids, first aggregates of the same experimental condition were pooled in plastic round-bottom wells. Then, all the gastruloids in a well were transferred to a single well of a glass-bottom plate (typically 10 gastruloids per well). This pooling increased the efficiency of locating gastruloids on the microscope. Gastruloids were typically imaged using a 10x air objective. Stitched images were acquired if the gastruloids did not fit in a single frame. For stained gastruloids, 100 μm confocal image stacks were collected, with 10 μm steps, encompassing the volume from the bottom of the gastruloid where it contacts the glass to approximately the axial midline.

To quantify fluorescence profiles along the gastruloid major axis, first a maximum-intensity z-projection was performed. Then, the gastruloid perimeter and major axis pixel coordinates were acquired in ImageJ, and these coordinates along with the max-intensity .tif file were imported into MATLAB (MathWorks). Here, custom code generates an arbitrary number (in this case, 200) minor axis lines that run perpendicular to the imported major axis, contained within the boundaries specified by the gastruloid perimeter. The average pixel intensity along each of these orthogonal lines is computed, thus providing a mean pixel intensity at discrete points along the major axis. These profiles can be averaged for multiple gastruloids of the same experimental condition to generate average plots with standard deviations, as seen throughout this paper. To detect ppErk peaks in an unbiased manner, we used the Matlab function ‘findpeaks’ to locate local maxima in the data. Note that, for all analyses, gastruloids were pooled and quantified without rejecting samples. Any difference between the number of gastruloids seeded and the number quantified per condition is a result of losses during the fixation and staining process. Starting numbers of gastruloids for each experiment are indicated in the figure captions. In the minor fraction of gastruloids with multiple posterior poles, the dominant pole was selected for quantification.

### Live imaging

All timelapse experiments were performed with gastruloids embedded in 50% Matrigel (Corning, 356231) / 50% N2B27 by volume to restrict translation of the gastruloid during imaging. The following protocol was adapted from^10^. To prepare for live imaging, first thaw Matrigel at 4°C overnight. Note: Matrigel must be kept cold to prevent polymerization and solidification. The following day prepare 50% Matrigel / 50% N2B27 solution in a 4°C room. Pipette this mixture into the center coverslip region of a 35 mm imaging dish (Ibidi, 81156). Taking a bucket filled with ice, place a flat metal object on the ice and put the imaging dish on top of the metal. This facilitates heat transfer, keeping the Matrigel cold. At a dissection microscope, pipette up individual gastruloids using a 200 μL pipette with the tip cut off. Deposit the desired amount of gastruloids into the Matrigel in the center of the imaging dish. If the gastruloids are clumped together they can be gently pipetted up from the Matrigel and distributed more evenly. Once this is complete, place the dish in the incubator at 37°C for 10 minutes, allowing the Matrigel to solidify. Then, add 2 mL of pre-warmed N2B27 to the dish.

Gastruloids were maintained at 37°C with 5% CO2 for the duration of all timelapse experiments using an environmental control unit (OKOLAB). The media in the imaging dish was covered with mineral oil (Sigma Aldrich, 330779) to prevent evaporation. The microscope was configured for differential interference contrast (DIC) microscopy for live imaging. Images were acquired every 2 minutes using a 20x air objective.

Particle image velocimetry (PIV) outputs were generated using the MATLAB application PIVlab^40^. Analysis was performed on 30 frames (1 hour) of data preceding the timestamp on the figure. Masks were drawn on the image to exclude background, and a fast-Fourier transform-based multipass algorithm was used to generate the velocity field for a given pair of frames. The mean velocity vector is displayed on the plot, and the heat map represents velocity magnitude at a given position.

### Gastruloid cell counting

A Countess II automated cell counter (Thermo Fisher) was used for all cell counting experiments. First, gastruloids were pooled into a single well of a round-bottom plastic 96-well plate. Pooling gastruloids provided a sufficient number of cells to obtain an accurate cell count. Between 5 and 10 gastruloids were pooled per measurement, depending on the experiment (see figure captions for experiment-specific values). All of the media was removed from the wells, and gastruloids were washed with 200 μL DPBS (-)Calcium (-)Magnesium, then transferred in the solution to a microcentrifuge tube. This was centrifuged at 400 relative centrifugal force for 5 minutes to pellet the gastruloids. The supernatant was then aspirated, and 100 μL TrypLE Express Enzyme was added to each tube, pipetting with sufficient force to disturb the gastruloid pellet. The samples were incubated in a 37°C bead bath for 5 minutes, then the solution was pipetted up and down to break gastruloids into a single-cell suspension. Next, 100 μL of 0.4% trypan blue solution (Gibco, 15250061) was added to each sample, and the samples were run on the automated cell counter to quantify total cell number and percent live/dead cells.

## Author Contributions

Conceptualization, E.J.U. and J.E.T.; Methodology, E.J.U. and J.E.T.; Investigation, E.J.U.; Resources, J.E.T.; Writing and Editing, E.J.U. and J.E.T.; Funding Acquisition and Supervision, J.E.T.

## Acknowledgements

The authors thank members of the Toettcher laboratory, particularly Harry McNamara, for insightful comments and suggestions. We also thank Alfonso Martinez Arias for generously sharing mESC lines, and gastruloid protocols. The project was supported by NSF CAREER Award 1750663 (J.E.T.) and NSF RECODE grant 2134935.

## Supplementary Figures

**Figure S1:**
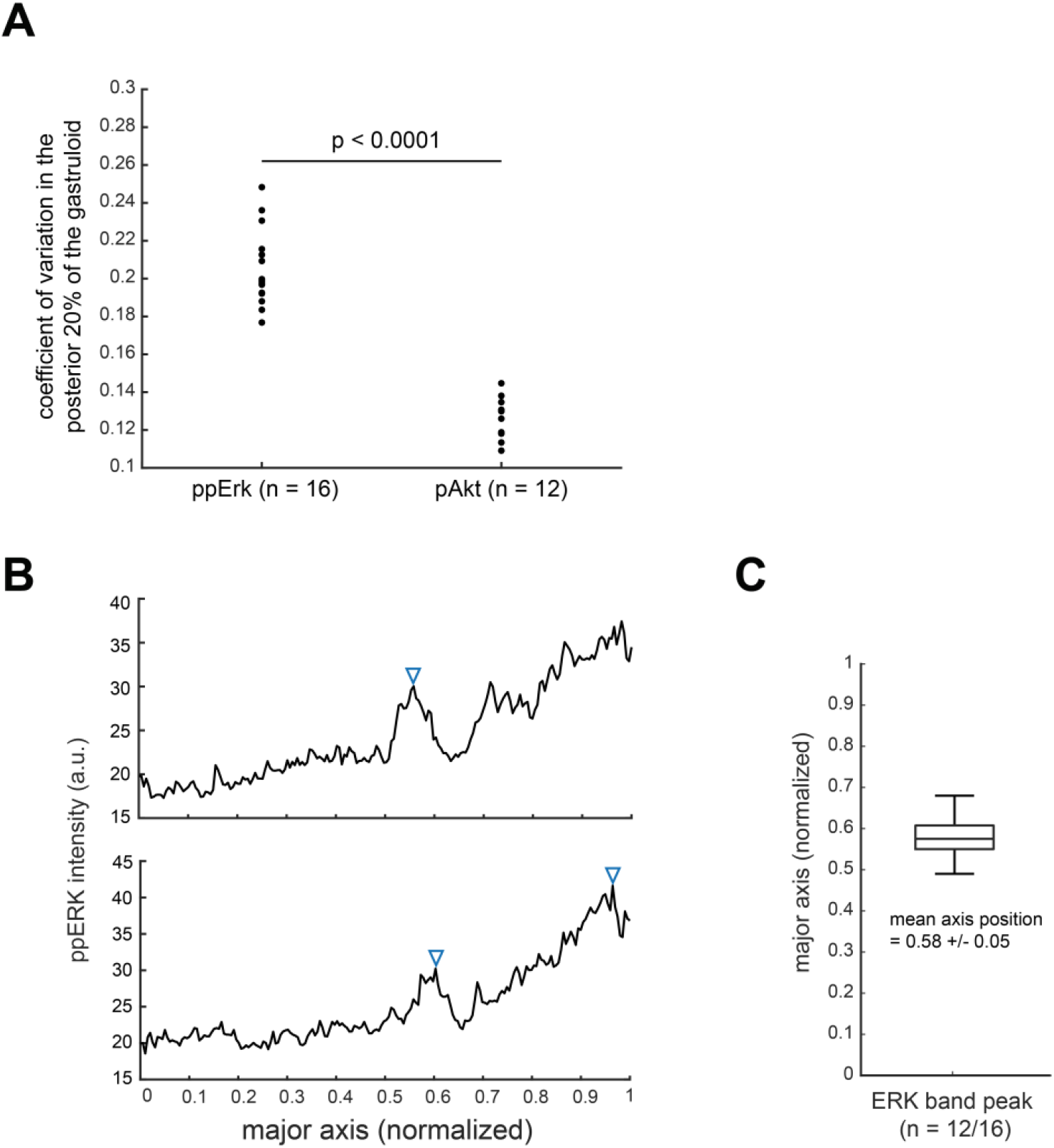
ppErk band detection. (A) The coefficient of variation (CV), calculated as the standard deviation divided by the mean, represents a measure of variability in a population. Here the CV is plotted for the fluorescence intensity in the posterior 20% of gastruloids that have been stained for either ppErk or pAkt. (B) Sample traces of ppErk intensity profiles with a secondary peak. MATLAB findpeaks() function was used to detect peaks in an unbiased manner. (top) Here, a peak was detected at an x position of 0.56, marked by a blue triangle. (bottom) If two peaks were detected, the anterior-most peak was used in subsequent analyses. (C) ppErk secondary peaks were detected in 12 of the 16 gastruloids analyzed. The peak positions are represented here in a box-and-whisker plot, with a mean axial position of 0.58 and a standard deviation of 0.05.

**Figure S2:**
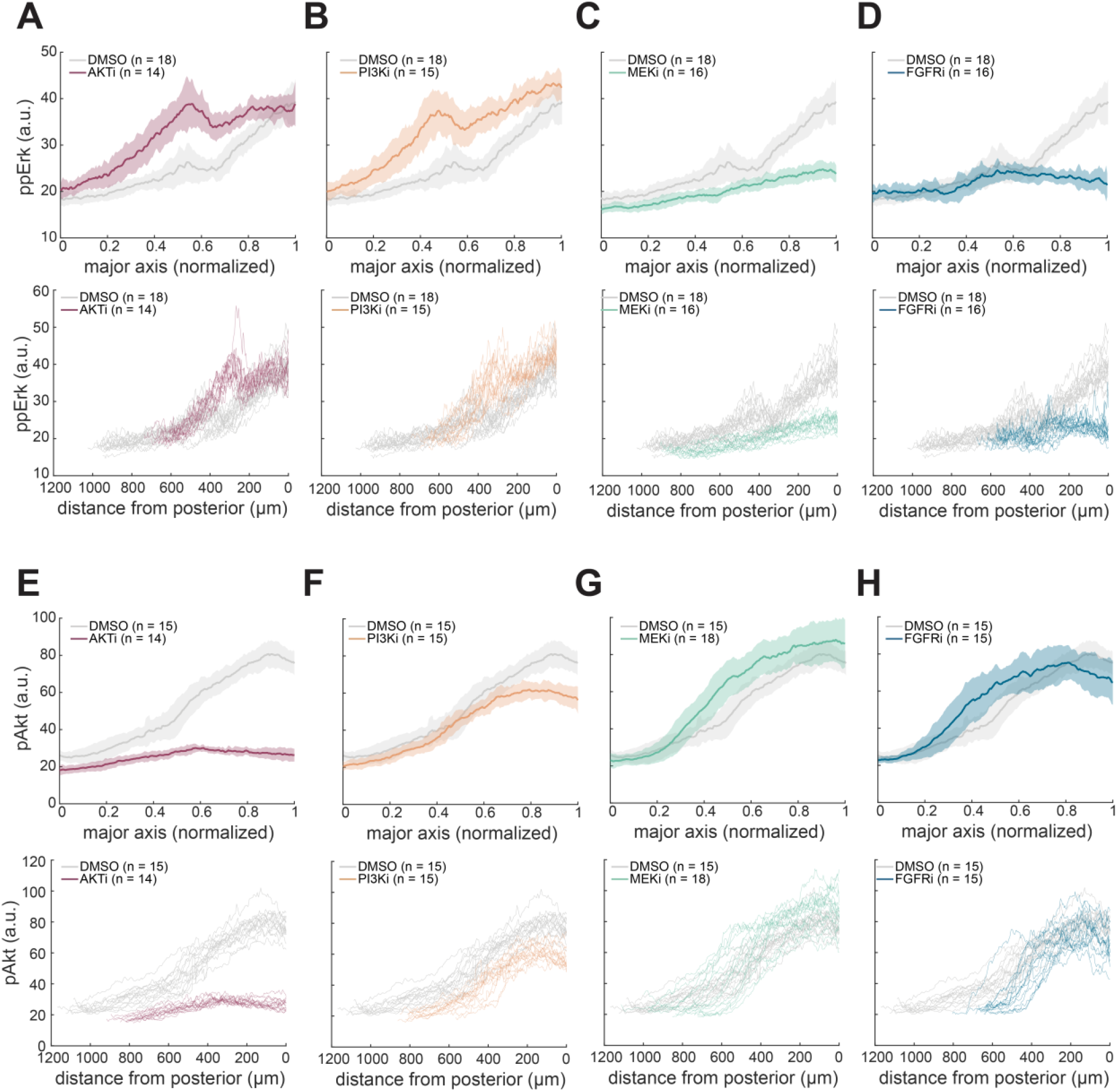
ppErk and pAkt gradients in response to inhibitor treatments. (A-D) ppErk spatial gradients for 96-120 h inhibitor-treated gastruloids compared to a DMSO control. (top) Mean curves and standard deviations are plotted as a function of normalized major axis position, where 0 is the anterior pole and 1 is the posterior pole. (bottom) Individual ppErk traces are plotted for the same data, here as a function of absolute distance from the posterior pole. (E-H) pAkt spatial gradients for 96-120 h inhibitor-treated gastruloids compared to a DMSO control. (top) Mean curves and standard deviations are plotted as a function of normalized major axis position, where 0 is the anterior pole and 1 is the posterior pole. (bottom) Individual pAkt traces are plotted for the same data, here as a function of absolute distance from the posterior pole. Each analysis began with 20 gastruloids. (N = 2 biological replicates).

**Figure S3:**
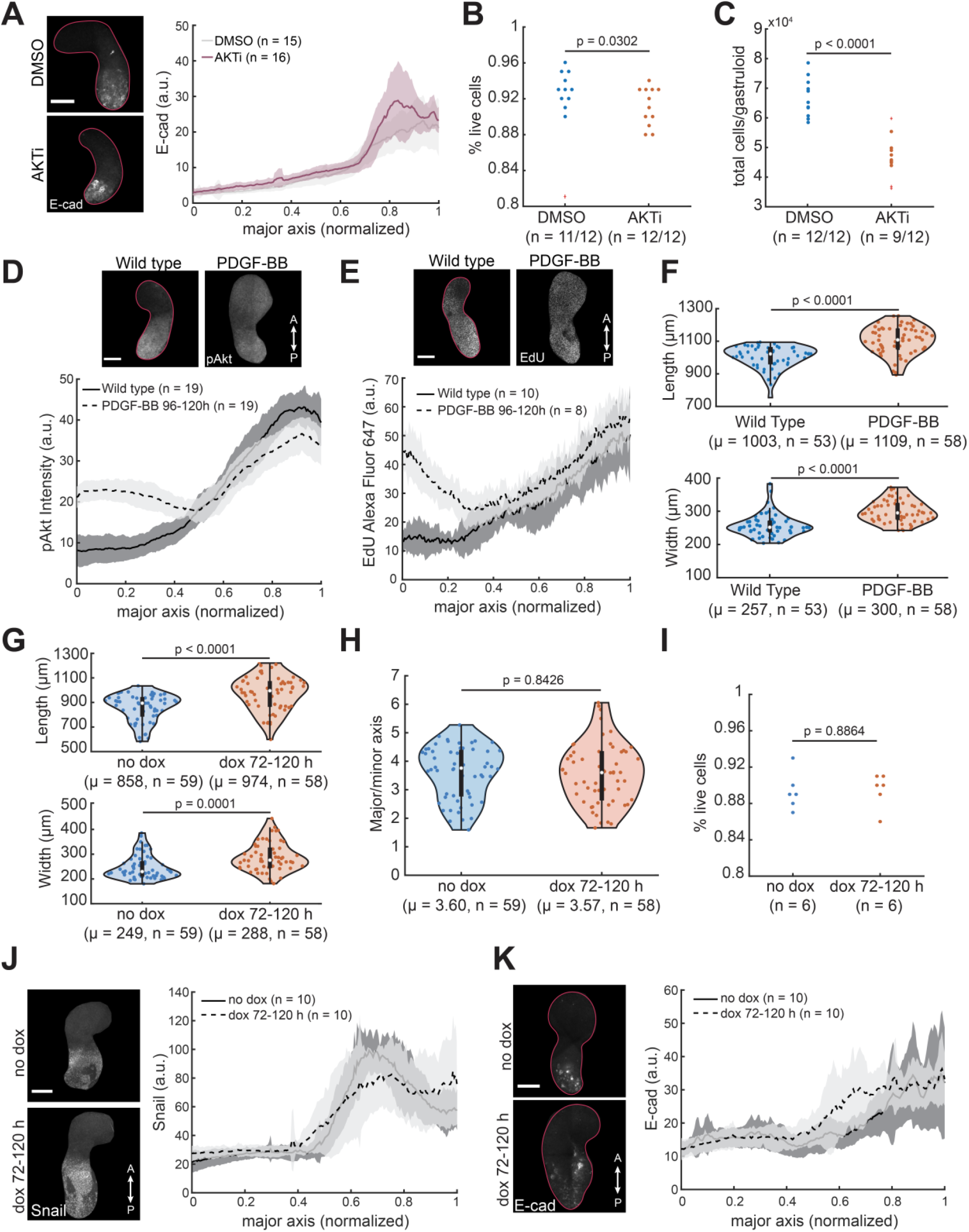
Consequences of Akt inhibition and hyperactivation. (A) Spatial profile of E-cad expression after Akt inhibition, compared to vehicle control. Analysis began with 20 gastruloids each (N = 2). (B-C) Percent live cells and mean total cell count per gastruloid upon Akt inhibition compared to DMSO control. Each data point represents the average cell count found from pooling 5 individual gastruloids. Data is aggregated from N = 3 biological replicates. Data marked as outliers are greater than 1.5 interquartile ranges above the upper quartile or below the lower quartile. (D) Spatial profiles of pAkt staining intensity after 100 ng/mL PDGF-BB treatment from 96-120 h, compared to a wild-type control. Each analysis began with 20 gastruloids. (N = 2) (E) Spatial profiles of EdU staining intensity after 100 ng/mL PDGF-BB treatment from 96-120 h, compared to a wild-type control. Each analysis began with 10 gastruloids. (N = 3) (F) Length and width of gastruloids treated with 100 ng/mL PDGF-BB from 96-120 h, compared to a wild-type control. μ is the data mean. Each analysis began with 60 gastruloids. N = 3 biological replicates. (G-H) Length, width, and major/minor axis aspect ratio of dox-Akt gastruloids treated with 1 μg/mL dox from 72-120 h, compared to a no-dox control. μ is the data mean. Each analysis began with 60 gastruloids. N = 3 biological replicates. (I) Percent live cells per dox-Akt gastruloid upon treatment with 1 μg/mL dox from 72-120 h, compared to a no-dox control. Each data point represents the average cell count found from pooling 10 individual gastruloids. Data is aggregated from N = 3 biological replicates. (J-K) Spatial profiles of Snail and E-cad in dox-Akt gastruloids treated with 1 μg/mL dox from 72-120 h, compared to a no-dox control. Each analysis began with 10 gastruloids (N = 1). All scale bars are 200 μm.

**Figure S4:**
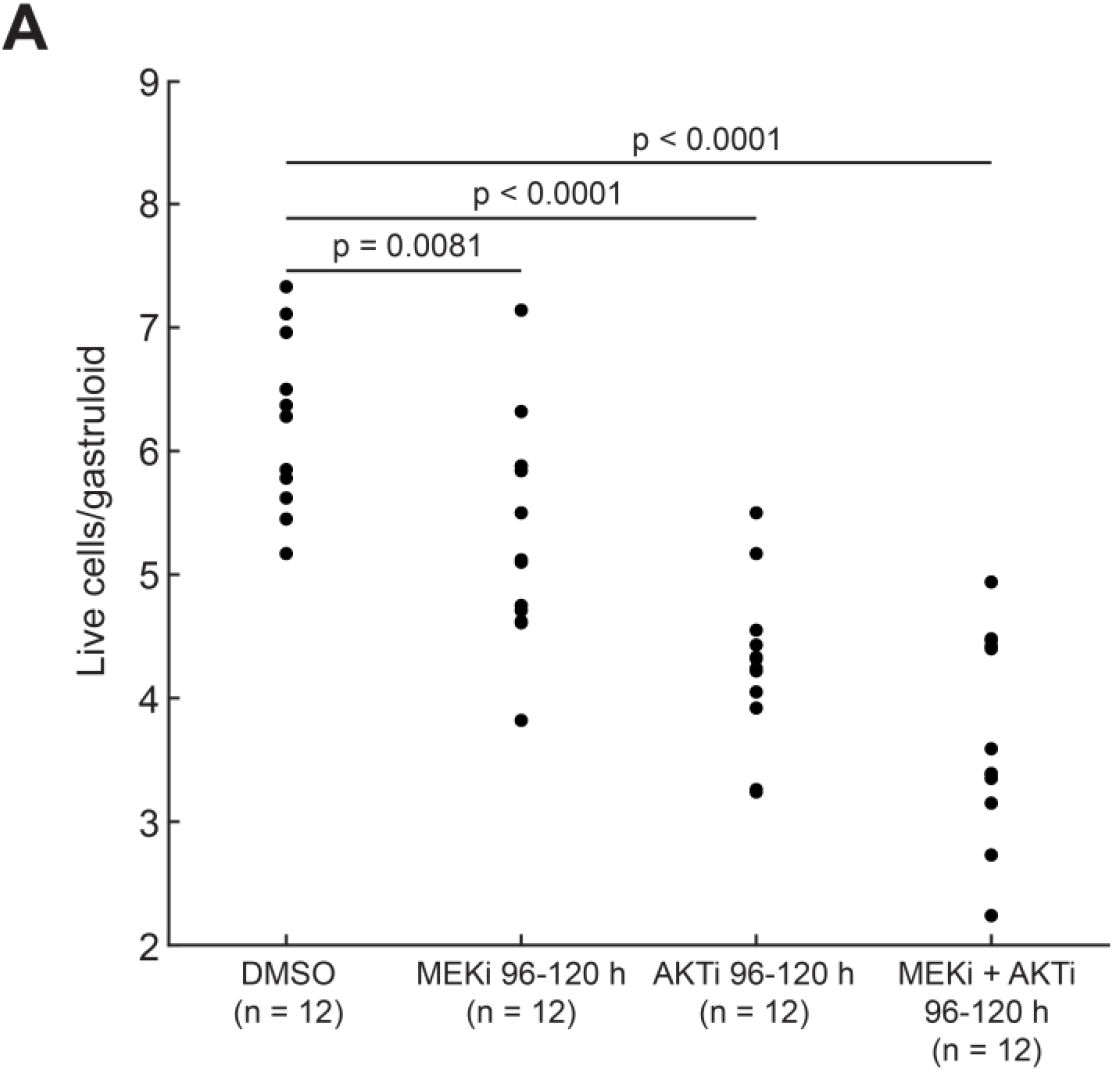
MEK inhibition results in a slight decrease in gastruloid total cell number. (A) Live cells per gastruloid upon MEK inhibition, Akt inhibition, or MEK + Akt inhibition from 96-120 h, compared to a DMSO control. Each data point represents the average cell count found from pooling 5 individual gastruloids. Data is aggregated from N = 3 biological replicates.

**Figure S5:**
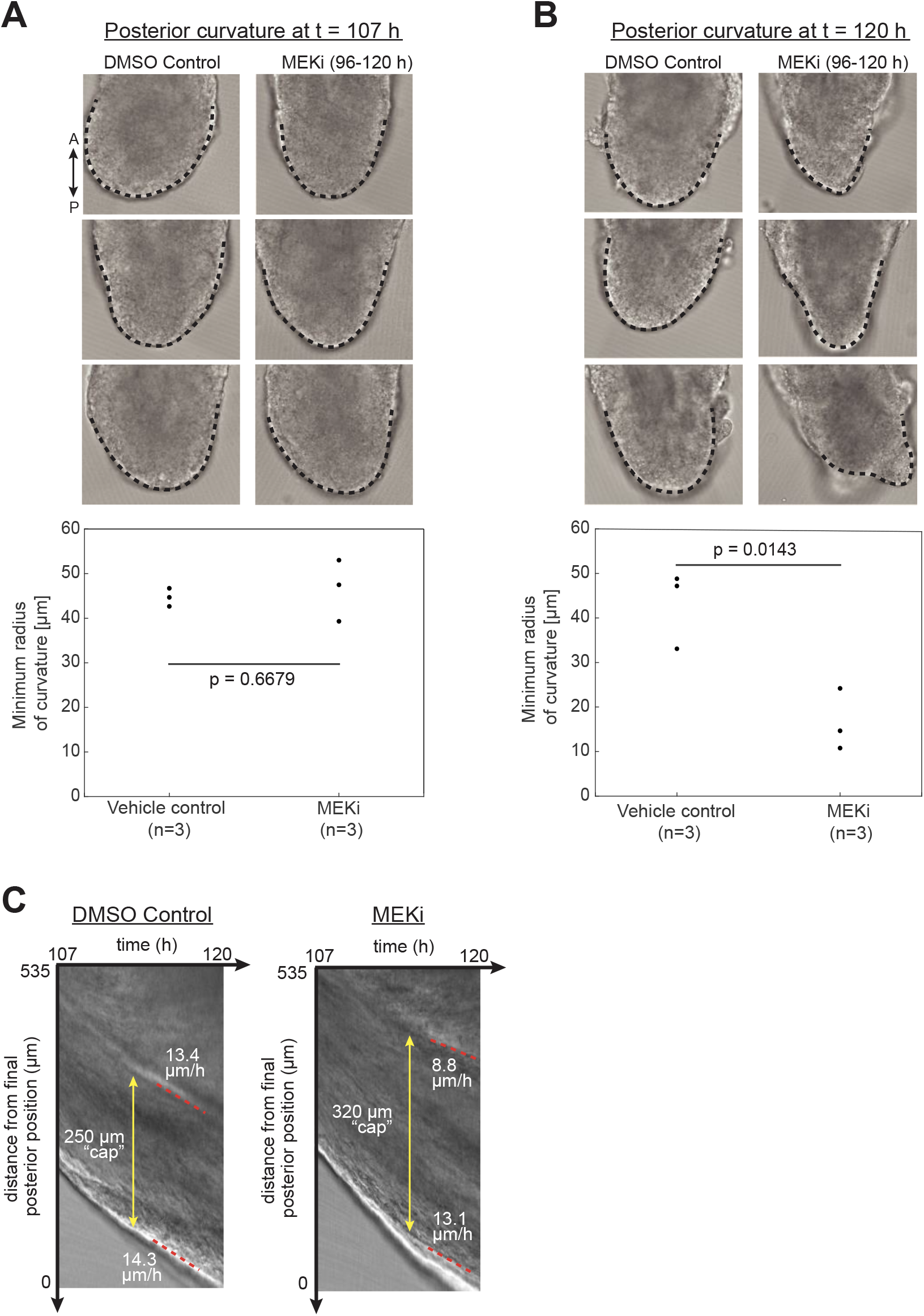
MEK Inhibition produces pointed posterior poles. (A-B, top) Frames from Movies 2 and 3 at either the beginning (107 h) or end (120 h) of the timelapse, demonstrating that MEK inhibited gastruloids grow a pointer posterior when compared to a control. Gastruloids were embedded in 50% Matrigel / 50% N2B27 over the course of imaging. The black dotted line traces the posterior pole of each gastruloid. (A-B, bottom) Quantification of minimum radius of curvature of the spline that traces the posterior pole for each gastruloid. (C) Kymographs generated from a vertical line along the gastruloid major axis for a MEKi gastruloid and DMSO control.

## Movie Legends

**Movie S1: Early gastruloid elongation.** Gastruloids were embedded in 50% Matrigel / 50% N2B27 in a 35 mm imaging dish at 95 h and imaged during the beginning of elongation (96-104 h). The sample was maintained at 37°C with 5% CO2 for the duration of the timelapse and imaged using differential interference contrast (DIC) microscopy with a 20x air objective. Images were acquired every two minutes. Posterior is oriented towards the bottom of the frame, as confirmed by endpoint Bra^GFP^ imaging.

**Movie S2: Late gastruloid elongation.** Equivalent experimental setup to Movie 1, except here gastruloids were embedded in 50% Matrigel / 50% N2B27 at 106 h and were imaged from 107-120 h. Images were acquired every two minutes. Posterior is oriented towards the bottom of the frame.

**Movie S3: Late gastruloid elongation in the presence of a MEK inhibitor.** Equivalent experimental setup to Movie 2, except here gastruloids were pre-treated in MEK inhibitor from 96 to 106 h, then were embedded in 50% Matrigel / 50% N2B27 at 106 h where they were again supplied with MEK inhibitor in the surrounding media. Gastruloids were live-imaged from 107-120 h, with images acquired every two minutes. Posterior is oriented towards the bottom of the frame. This timelapse was acquired simultaneously with Movie 2.

